# Resolving an unconventional non-photochemical quenching signature at the light-to-dark transition

**DOI:** 10.1101/2024.10.17.618902

**Authors:** Lam Lam, Dhruv Patel-Tupper, Henry E. Lam, Collin J. Steen, Alexa Ma, Sophia A. Ma, Anna Leipertz, Tsung-Yen Lee, Graham R. Fleming, Krishna K. Niyogi

## Abstract

Non-photochemical quenching (NPQ) protects photosynthetic organisms via diverse molecular players contributing at varying timescales. However, in the absence of one of the largest contributors to NPQ, energy-dependent quenching (qE), we observe an unusual but universal phenomenon: a transient increase in quenching in the dark following high light exposure. To mechanistically interrogate this light-to-dark (LtD) NPQ phenotype, we performed chlorophyll fluorescence lifetime snapshot measurements across a diverse array of Arabidopsis mutant backgrounds and chemical treatments. We found that the electrochemical gradient across the thylakoid membrane is essential for this phenomenon. Through analysis of higher-order Arabidopsis mutants, we also found that LtD NPQ is independent of the known forms of photoprotective NPQ, as well as the major and minor light-harvesting complexes (LHCII). Our results point to LtD NPQ as a photoinhibition (qI)-related, reaction center quenching with implications for photoprotection in fluctuating light.

## INTRODUCTION

Photosynthesis in plants begins with the singlet-state excitation of chlorophyll (^1^Chl*) in light-harvesting complexes (LHCs), whose energy is eventually transferred to the reaction center (RC) where charge separation occurs. However, the capacity of photosynthesis to use absorbed light energy is not unlimited. Under high light, excess excitation energy can result in the formation of the longer-lived triplet state of chlorophyll (^3^Chl*) by intersystem crossing, which can produce reactive oxygen species (ROS) capable of photo-oxidative damage to the pigments, proteins, and lipids within the thylakoid membrane (Niyogi, 1999). Oxygenic photosynthetic organisms have evolved sophisticated photoprotection mechanisms, a series of non-photochemical quenching (NPQ) processes, that safely dissipate excess excitation energy as heat (Bassi and Dall’Osto, 2021).

Forward genetic screens have helped to increase our understanding of land plant NPQ. Functional NPQ requires photosystem II subunit S (PsbS/NPQ4) *in vivo* (Li et al., 2000), is augmented by zeaxanthin (Zea) produced by violaxanthin de-epoxidase (VDE/NPQ1), and is relaxed alongside the conversion of Zea back to violaxanthin (Vio) by zeaxanthin epoxidase (ZEP/ABA1/NPQ2) under light-limiting conditions (Niyogi et al., 1998). In *Arabidopsis thaliana*, lycopene epsilon cyclase (LUT2) is essential for the production of the carotenoid lutein (Lut), which additively contributes to NPQ independently of Zea (Pogson et al., 1996; Pogson et al., 1998). In the absence of both Zea and Lut (i.e., the *npq1 lut2* mutant), PsbS cannot induce rapidly reversible NPQ (Niyogi et al., 2001).

Cumulatively, PsbS, Zea, and Lut are responsible for seconds-minutes timescale energy-dependent quenching (qE). A longer-acting, Zea-dependent quenching (qZ), contributes independently of the ΔpH necessary to trigger qE (Nilkens et al., 2010). Unlike qE and qZ, LHCII state transitions (qT) do not directly quench ^1^Chl*, but do regulate light absorption by partitioning excitation between photosystem I (PSI) and PSII (Bellafiore et al., 2005). A recently discovered form of sustained antenna quenching (qH) has been found to be chloroplast lipocalin-dependent, and occurs under stress conditions such as cold and high light (Malnoë et al., 2018). The most slowly relaxing component of NPQ, photoinhibitory quenching (qI), results from photoinactivation of PSII by ROS—relaxing on the timescale of hours to days (Long et al., 1994). In concert, NPQ mechanisms involve different molecular players to sustain plant fitness at varying light intensities and timescales.

qE accounts for the vast majority of NPQ induction and relaxation in response to high light (HL) and darkness (D), respectively, under most conditions. However, in the absence of qE in plants (Dall’Osto et al., 2014a; Steen et al., 2020), green algae (Steen et al., 2022), stramenopile algae (Perin et al., 2023), and even diatoms (Buck et al., 2019), we observe a conserved and unusual phenomenon: a rapid, transient increase in NPQ in the dark, after the transition from HL. The source of this non-canonical behavior, hereafter termed light-to-dark transition-dependent NPQ, or LtD NPQ, has not yet been investigated. A better understanding of its possible contributions to wild-type (WT) and qE-less NPQ may expand our understanding of photoprotection in fluctuating light environments.

To this end, we carried out a systematic investigation of the LtD NPQ response via a comprehensive array of higher-order *A. thaliana npq4* mutants (**Table 1**) and chemical inhibitors (**Table 2**) to interrogate the factors necessary for LtD NPQ. We measured changes in the fluorescence lifetime of ^1^Chl* in intact leaves during a periodically fluctuating actinic light sequence by time-correlated single photon counting (Sylak-Glassman et al., 2016), to quantify the capacity of quenching and its kinetics without the influence of chloroplast movement, enhanced light scattering, or Chl concentration differences (see Materials and Methods). Leveraging a process-of-elimination approach across genetic backgrounds and conditions, we hypothesize that LtD NPQ is a qI-type reaction center quenching that manifests in the absence of major NPQ components such as qE. Further work to resolve the phenotypic consequences of LtD NPQ will expand our understanding of NPQ in dynamic light environments.

**Table 1.**
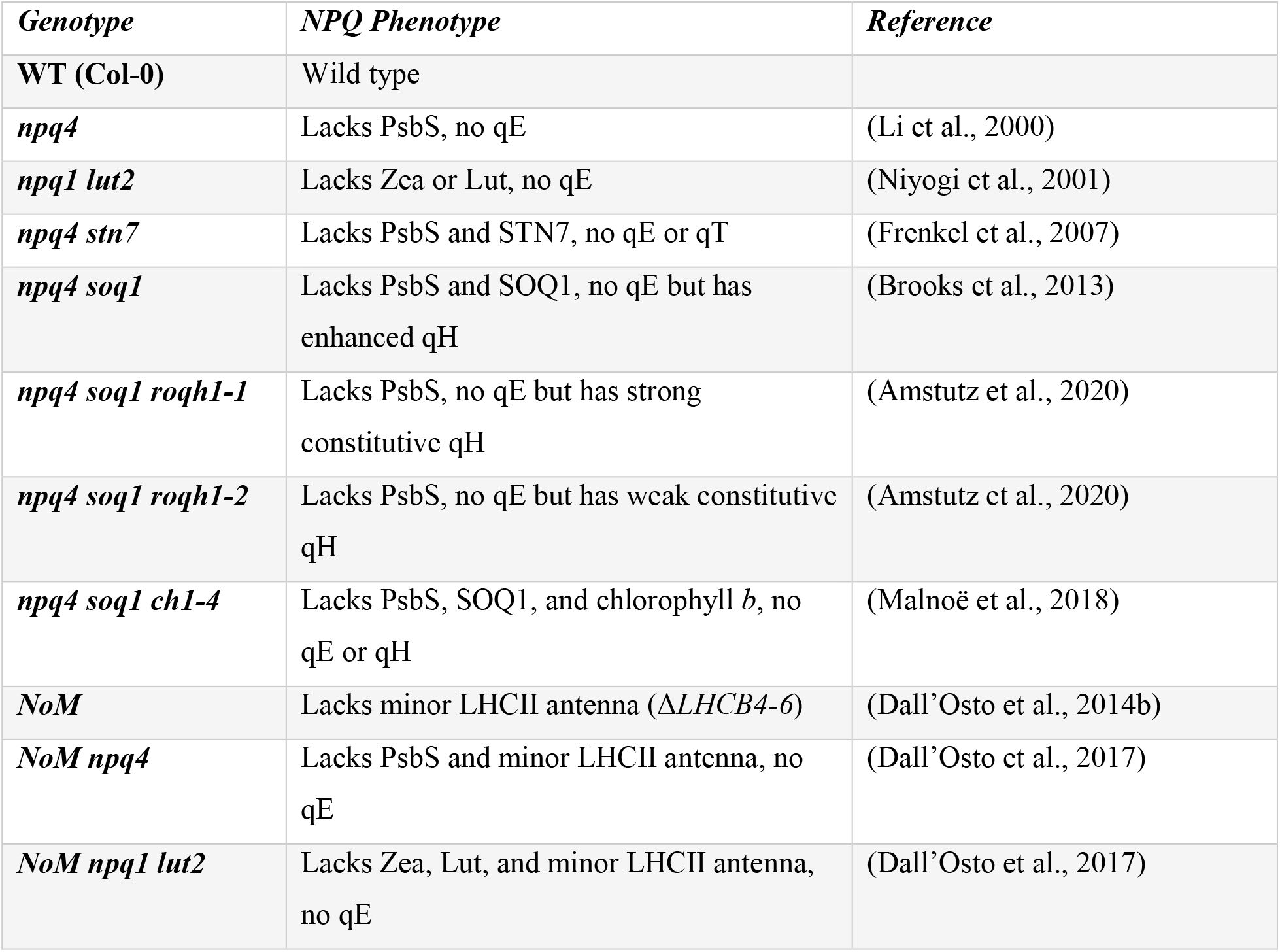
Summary of Arabidopsis lines used in this study.

**Table 2.**
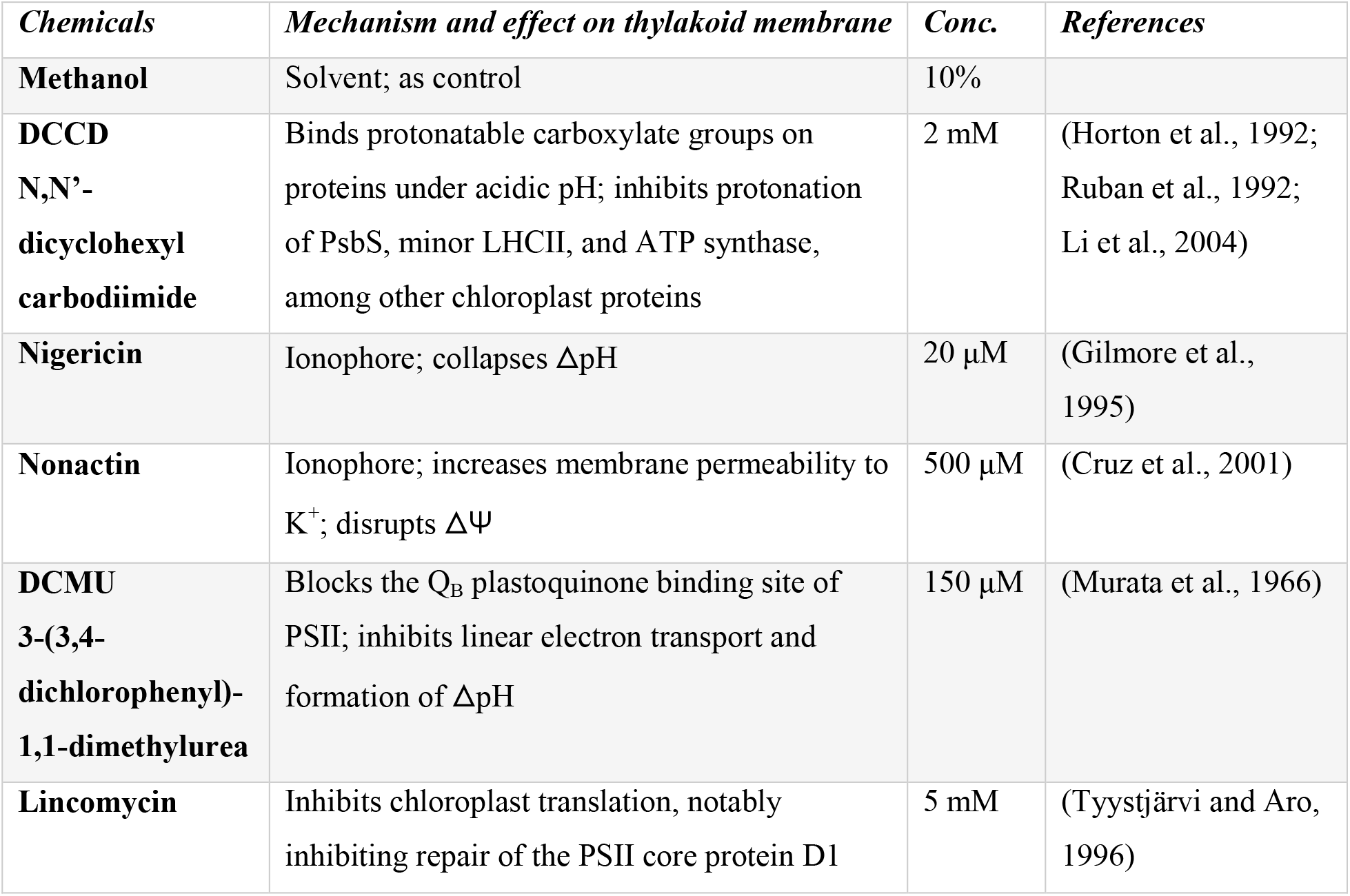
Summary of chemical inhibitors and working concentrations used in this study.

| *Chemicals* | *Mechanism and effect on thylakoid membrane* | *Conc.* | *References* |
| --- | --- | --- | --- |
| **Methanol** | Solvent; as control | 10% |  |
| **DCCD**<br>**N,N'-**<br>**dicyclohexyl**<br>**carbodiimide** | Binds protonatable carboxylate groups on proteins under acidic pH; inhibits protonation of PsbS, minor LHCII, and ATP synthase, among other chloroplast proteins | 2 mM | (Horton et al., 1992; Ruban et al., 1992; Li et al., 2004) |
| **Nigericin** | Ionophore; collapses ΔpH | 20 μM | (Gilmore et al., 1995) |
| **Nonactin** | Ionophore; increases membrane permeability to $K^+$; disrupts $\Delta\Psi$ | 500 μM | (Cruz et al., 2001) |
| **DCMU**<br>**3-(3,4-**<br>**dichlorophenyl)-**<br>**1,1-dimethylurea** | Blocks the $Q_B$ plastoquinone binding site of PSII; inhibits linear electron transport and formation of ΔpH | 150 μM | (Murata et al., 1966) |
| **Lincomycin** | Inhibits chloroplast translation, notably inhibiting repair of the PSII core protein D1 | 5 mM | (Tyystjärvi and Aro, 1996) |

## MATERIALS AND METHODS

### Plant Genotypes and Growth Conditions

WT (Col-0 ecotype) and the 10 *Arabidopsis thaliana* NPQ mutant lines used in this study are summarized in **Table 1**. Plants were germinated on ½ MS plates and then transplanted to soil (Sunshine Mix #1, Sungro) after 2 weeks. Plants were grown under 110 μmol photons m^−2^ s^−1^ on a 10-h day and 14-h night cycle in a growth chamber set at 23 °C. 5-6 week old plants were used for measurements.

### Syringe Infiltration with Chemicals

Plants were dark-acclimated for 1 hour before infiltration. Healthy, mature leaves were gently infiltrated by syringe with chemicals at the concentrations listed in **Table 2**. All chemicals were solubilized in either 100% methanol or water and diluted with distilled deionized water before each infiltration. The final concentrations of methanol did not exceed 10% (v/v). After infiltration, each leaf remained on the plant and was dark-acclimated for another 25 min before fluorescence lifetime measurements. For experiments involving high light (HL) pre-treatment, leaves were infiltrated and whole plants were exposed to 400 μmol photons m^−2^ s^−1^ for 16 h, followed by a 1 h dark-acclimation before time-correlated single photon counting.

### Fluorescence Lifetime Measurements and Analysis

To measure Chl fluorescence lifetimes and quantify the quenching capacity and kinetics of the leaf samples, we applied a custom-built time-correlated single photon counting (TCSPC) setup, as previously described (Steen et al., 2020). Pulses centered at 808 nm were generated by a Ti:sapphire oscillator (Coherent, Mira900f, 76 MHz) pumped by a Coherent Verdi G10 diode laser with its center wavelength set at ∼532 nm. The frequency was doubled to ∼404 nm after passing through a beta-barium borate crystal to excite the Chl *a* Q_x_ band. A portion of the excitation beam was divided by a beam splitter to a sync photodiode (Becker-Hickl, PHD-400), providing time references to the TCSPC card. The remainder of the excitation beam was then incident at an ∼70° angle to the adaxial side of the leaf while avoiding the leaf stem. For infiltrated leaf samples, the beam was directed on the infiltrated region, avoiding any mechanical lesion caused by the syringe infiltration. Leaves were dark-acclimated for 1 h before measurements. Reaction centers were closed by the excitation beam with its power set to 1.0 mW. Leaves were exposed to a 40 min actinic light (Leica KL1500 LCD) sequence, composed of 2 cycles of 10 min high light (600 µmol photons m^-2^ s^-1^) and 10 min dark at 15 s measurement resolution. A microchannel plate (MCP)-photomultiplier tube (PMT) detector (Hamamatsu R3809U MCP-PMT) was used after a monochromator (HORIBA Jobin-Yvon; H-20), which was set to 680 nm to exclusively collect the Chl *a* Qy band fluorescence. The FWHM of the instrument response function was 36−38 ps, sufficient to resolve Chl fluorescence lifetimes on the sub-ns timescale. A LabVIEW program controlled a series of shutters, which coordinated the sequencing of the excitation beam, actinic light, and detector.

Within a 1 s total integration time of detection, a 0.2 s timestep of the data showing the longest lifetime was selected for further data processing to ensure that the reaction centers were closed (Sylak-Glassman et al., 2016). Each fluorescence decay profile was fitted with a bi-exponential decay function and the amplitude-weighted average lifetime 𝜏 was calculated by:

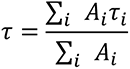

where A_i_ and τ_i_ are the amplitudes and fluorescence lifetimes of the ith fitting component, respectively. The NPQ capacity is defined by

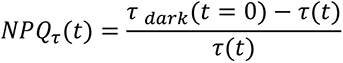

where *τ_dark_ (t=0)* is the average of amplitude-weighted average lifetimes of the three initial dark snapshots, and τ(t) is amplitude-weighted average lifetime at the corresponding snapshot sequence time *t*.

### SDS-PAGE and immunoblotting

For isolation of total protein, infiltrated portions of leaves were excised, ground in a solution of 5 mM EDTA pH 8.0, 20 mM HEPES pH 7.6, 5% (w/v) LDS, 60 mM DTT, and incubated in the dark at room temperature for 30 min. The proteinaceous supernatant was collected after centrifugation at 14,000 rpm at 4 °C for 10 min. Total chlorophyll content of the supernatant was quantified as described (Porra et al., 1989). 1 µg of total chlorophyll was loaded per well for SDS-PAGE.

Solubilized proteins were separated by size using BIO-RAD Any kD™ Mini-PROTEAN TGX Stain- Free™ Protein gels. Coomassie staining was performed by incubating the gel in fixing solution (50% Methanol, 5% Acetic acid) for 20 min, followed by 15 min incubation in 2.5g/L CBB G-250 solution (55% Methanol 5% Acetic acid), and overnight destaining (5% Methanol, 7.5% Acetic acid) before imaging with the BIO-RAD ChemiDoc MP imaging system. A duplicate set of gels were transferred to a polyvinylidene difluoride membrane (Immobilon-FL 0.45 μm, Millipore) via semi-dry transfer using the BIO-RAD Trans-Blot Turbo Blotting Instrument. Polyclonal antibodies raised against the D1 DE-loop (AS10 704) were obtained from Agrisera (Sweden) and used for immunoblotting. Donkey anti-rabbit IgG antibody with horseradish peroxidase (1:10,000; GE Healthcare) was used as a secondary antibody and visualized using the SuperSignal West Femto Maximum Sensitivity Substrate (Thermo Scientific) chemiluminescent system on the BIO-RAD ChemiDoc MP imaging system.

## RESULTS

### Non-canonical NPQ in the absence of qE

We utilized two light-dark cycles to resolve the transient increase in NPQ observed after actinic light transitions, termed light-to-dark transition-dependent (LtD) NPQ, by time-correlated single photon counting (TCSPC). LtD NPQ was observed in Arabidopsis *npq4* leaves (lacking PsbS and qE), but not in WT leaves after both light-to-dark transition points (t = 10 min, 30 min), as denoted by a transient decrease in fluorescence lifetime (**Fig. 1a**) and corresponding increase in NPQτ (**Fig. 1b**). Analysis of the stepwise change in average fluorescence lifetimes between timepoints highlighted the substantial drop in *npq4* fluorescence lifetime after the transition to darkness. In contrast, WT exhibited rapid relaxation of NPQ, as evidenced by the steady rate of WT fluorescence lifetime recovery (0.02-0.05 ns / 15 s interval) after both transition points (**Fig. 1c**).

**Figure 1.**
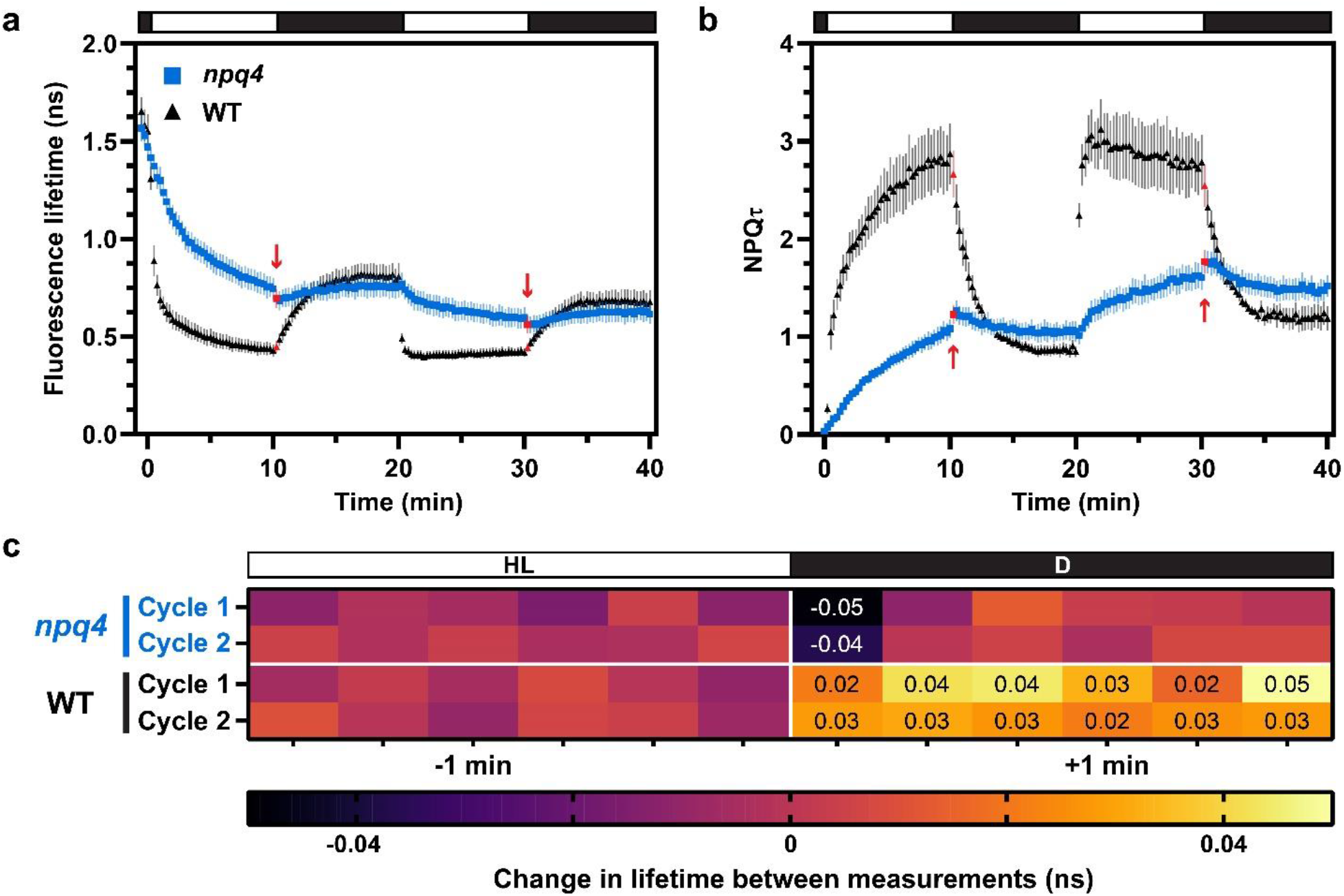
*Changes in WT and npq4 fluorescence lifetime and NPQτ values in a periodic light/dark sequence*. **(a)** Fluorescence lifetimes and **(b)** NPQτ for wild-type (black, n=4) and *npq4* (blue, n=7) leaves in response to two cycles of 10 min high light (600 µmol photons m^-2^ s^-1^) and 10 min darkness (0 µmol photons m^-2^ s^-1^) at 15 s measurement intervals. Red arrows and symbols indicate the first 15 s dark timepoint following each light-to-dark transition. Error bars show ±1 SEM. **(c)** Heatmap of the average stepwise change in chlorophyll fluorescence lifetime between each 15 s interval, for measurements flanking each light-dark transition. Quenching of fluorescence lifetime is shown in black and purple, whereas fluorescence lifetime recovery is shown in yellow and orange. Values larger than ±0.02 ns are shown.

### Membrane energetics affects lifetime kinetics and LtD NPQ

To determine what native conditions support formation of LtD NPQ, we used various chemical inhibitors that differentially affect membrane energization (**Table 2**). All selected chemicals had a significant effect on WT NPQ (**Fig. 2a**). Whole-leaf infiltration of DCCD (disrupting protein protonation) did not slow NPQ induction, but significantly inhibited NPQ relaxation. In contrast both nigericin (collapsing the ΔpH) and nonactin (disrupting the ΔΨ) inhibited NPQ capacity. DCMU (inhibiting linear electron transport) had the strongest effect, significantly reducing NPQ induction and entirely abolishing relaxation.

**Figure 2.**
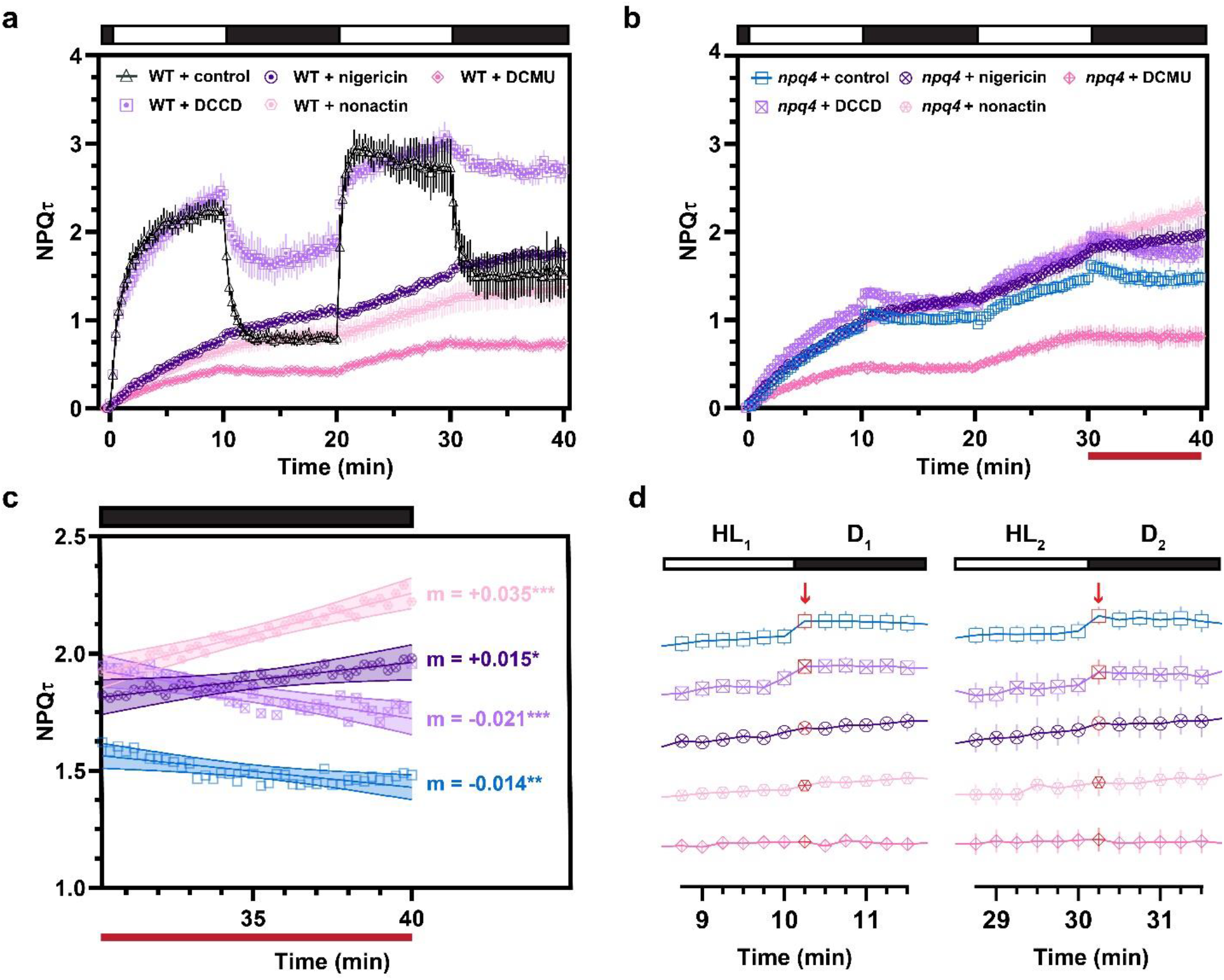
*NPQτ values for chemically inhibited WT and npq4 leaves*. **(a)** NPQτ of WT leaves infiltrated with either 10% methanol as control (black, n=3), 2 mM DCCD (purple, n=3), 20 µM nigericin (indigo, n=3), 500 µM nonactin (pink, n=4), or 150 µM DCMU (magenta, n=5). **(b)** NPQτ of *npq4* leaves infiltrated with either 10% methanol as control (blue, n=5), 2 mM DCCD (purple, n=4), 20 µM nigericin (indigo, n=4), 500 µM nonactin (pink, n=4), or 150 µM DCMU (magenta, n=5). **(c)** Linear regression of the 30 - 40 min dark relaxation from (b, red line) with asterisks signifying significantly non-zero slopes (* p < 0.5, ** p < 0.01, *** p < 0.001). **(d)** Arbitrarily staggered NPQτ values of *npq4* infiltrated leaves at each light-dark transition. Red arrows and symbols indicate the first 15 s timepoint after the light-to-dark transition.

When nigericin and nonactin were applied to *npq4* leaves, the kinetics of NPQ, but not its overall magnitude, were significantly affected (**Fig. 2b**). These differences were most pronounced in the second dark relaxation phase, where nigericin- and nonactin-treated leaves continued to slowly induce NPQ in the dark (increasing NPQτ values that resulted in a positive slope during relaxation), while NPQ in DCCD-treated leaves relaxed similarly to the 10% methanol control treatment (decreasing NPQτ values that resulted in a negative slope during relaxation) (**Fig. 2c**). The differences in relaxation after addition of nigericin and nonactin were consistent with the kinetics of infiltrated WT leaves shown in Fig. 2a. DCMU treatment in *npq4* entirely phenocopies the WT + DCMU phenotype (**Fig. 2a, 2b**). Nigericin, nonactin, and DCMU all appeared to reduce the formation of LtD NPQ at each light-to-dark transition (**Fig. 2d**), suggesting a role for membrane energetics in the LtD NPQ phenotype.

### The LtD NPQ response is independent of qE, qZ, qT, and qH

To study whether LtD NPQ arises from an NPQ mechanism other than qE, we screened various higher order qZ, qT, and qH NPQ mutants which also lacked qE under periodic light. Mutants lacking the NPQ- associated carotenoids Zea and Lut (*npq1 lut2*) or lacking PsbS and the STN7 kinase necessary for state transitions (*npq4 stn7*) still exhibited the *npq4*-like LtD NPQ phenotype (**Fig. 3a**). Previous studies have shown that LtD NPQ behavior still exists in the qE- and qH-deficient *npq4 soq1 lcnp* mutant (Malnoë et al., 2018). To further assess whether the LtD NPQ response was influenced by qH, we tested the constitutively qH-quenched *npq4 soq1 roqh1* mutants. However, even with moderate (*npq4 soq1 roqh1- 2*) or strong (*npq4 soq1 roqh1-1*) constitutive qH, LtD NPQ was still present (**Fig. 3b**) despite significant quenching of initial fluorescence lifetimes in the dark (**Fig. 3c**).

**Figure 3.**
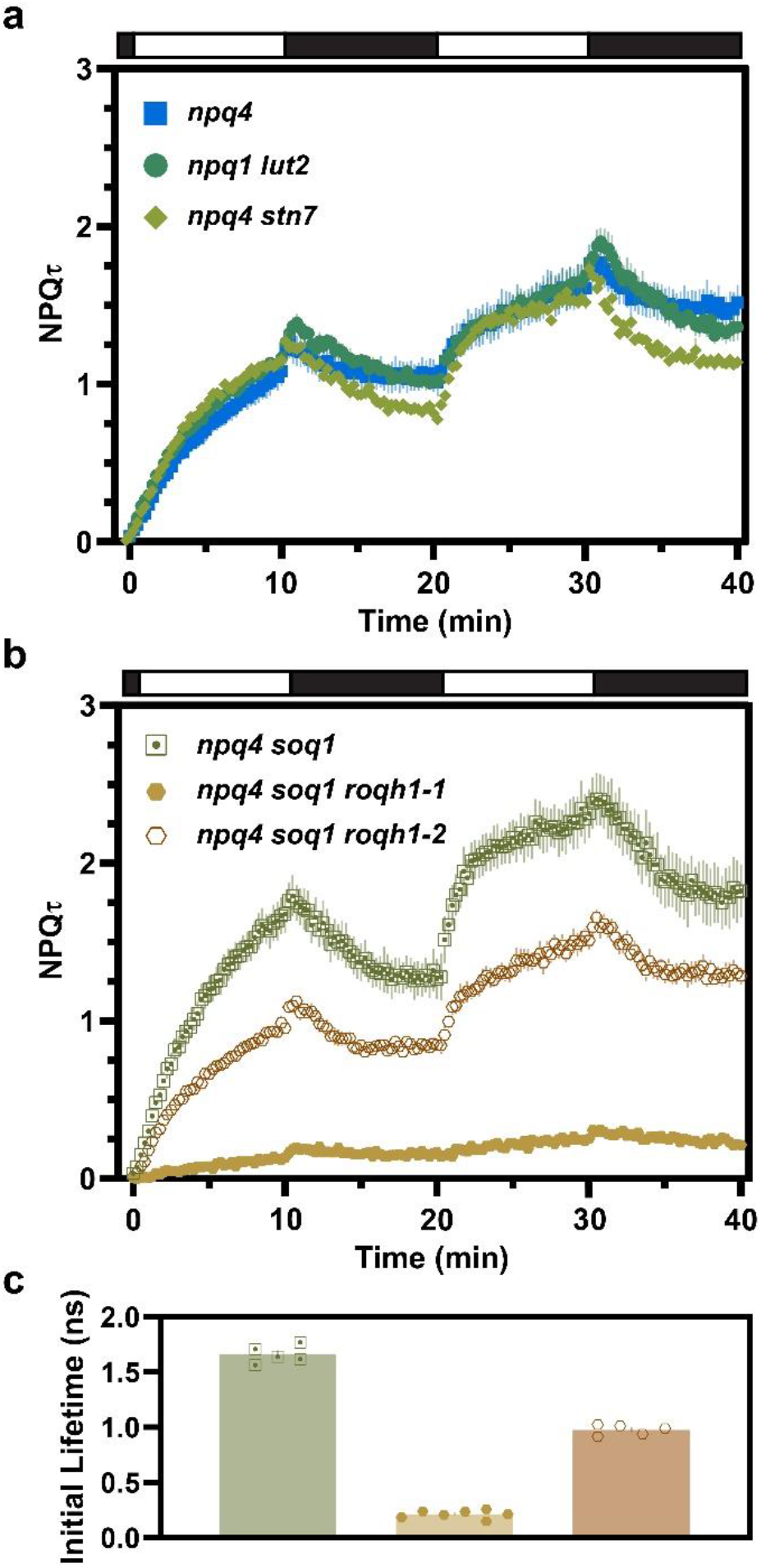
*NPQτ values for higher order npq4 mutants deficient in other known types of photoprotective NPQ*. **(a)** NPQτ of *npq1 lut2* (teal, n=5) and *npq4 stn7* (light green, n=3) compared to *npq4* (blue, n=7). **(b)** NPQτ of *npq4 soq1* (green, n=5) compared to *npq4 soq1 roqh1-1* (light brown, n=7) or *npq4 soq1 roqh1-2* (dark brown, n=5) alleles. **(c)** Dark-acclimated initial fluorescence lifetimes of *npq4 soq1 and npq4 soq1 roqh1* mutant lines. Symbols are the same as in panel b. Error bars show ±1 SEM.

### Major and minor light-harvesting antennae are not quenching sites for LtD NPQ

We next sought to clarify possible quenching sites responsible for LtD NPQ. We measured fluorescence lifetimes of a higher order *npq4* mutant lacking chlorophyll *b* (*chlorina1* or *ch1*), disrupting the stability of all LHCII proteins excluding LHCB5 (Murray and Kohorn, 1991; Espineda et al., 1999). Loss of chlorophyll *b* decreased the observed dark-acclimated fluorescence lifetime (**Fig. 4a**, inset) and nullified the phenotypic contribution of *soq1*-dependent NPQ, as previously reported (Malnoë et al., 2018).

**Figure 4.**
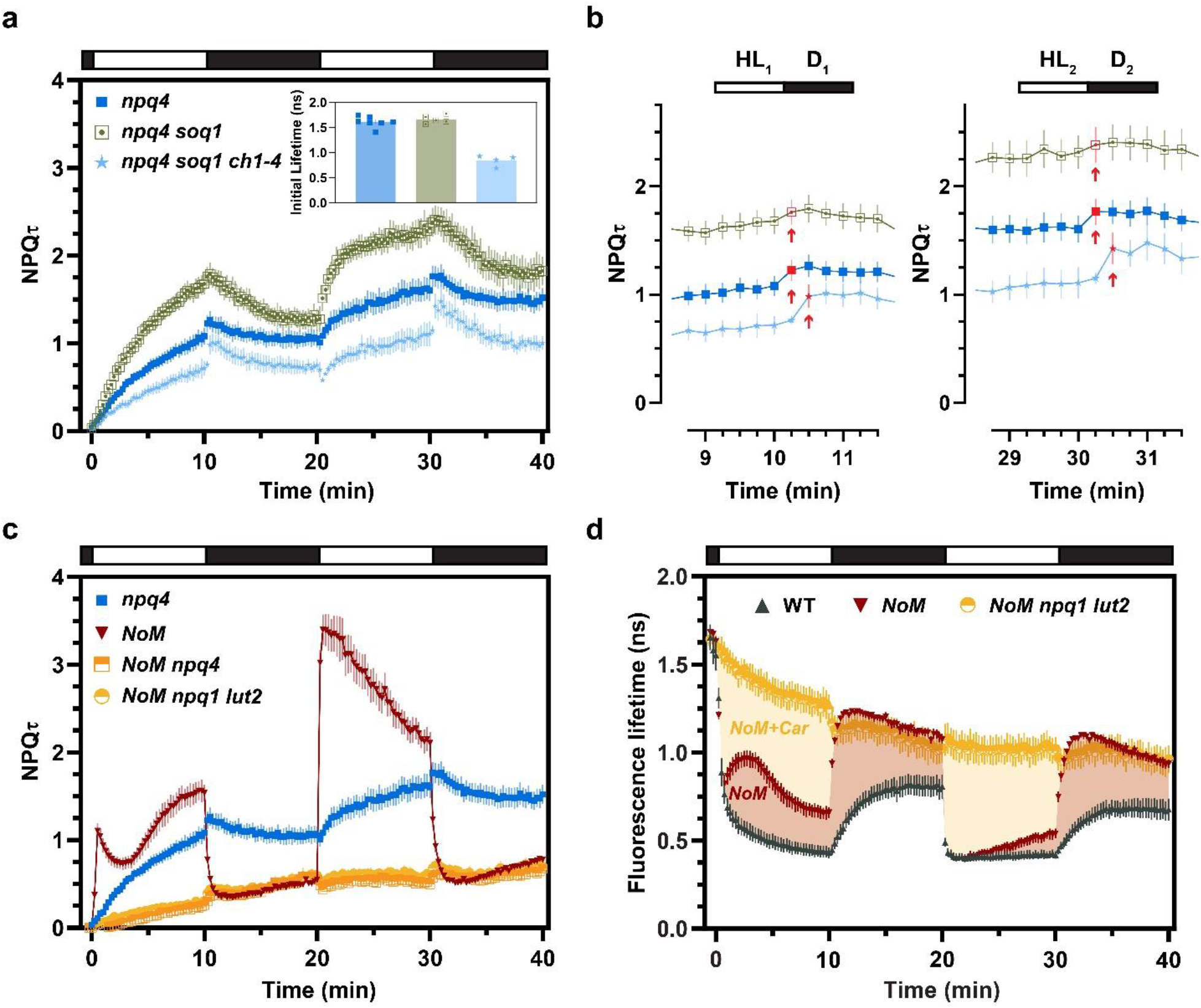
*NPQτ and fluorescence lifetime values of mutants lacking stable major and minor LHCII antenna*. **(a)** NPQτ of *npq4* (blue, n=7), *npq4 soq1* (green, n=5), and *npq4 soq1 ch1-4* (light blue, n=6). Inset in (a) shows differences in initial fluorescence lifetime. **(b)** NPQτ of sub-panel (a) at each light-dark transition. Red arrows and symbols denote the first timepoint with detectable LtD NPQ. (**c)** NPQτ of *npq4* (blue, n=7), *NoM* (red, n=5), and higher order *NoM* mutants lacking PsbS (*NoM npq4*, orange, n=5) or Zea and Lut (*NoM npq1 lut2*, yellow, n=4). **(d)** Fluorescence lifetimes of *NoM* and *NoM npq1 lut2* relative to WT (black, n=4). Error bars show ±1 SEM.

Despite this overall disruption we still observed the LtD NPQ response in *ch1* mutants (**Fig. 4a**). Surprisingly, the *ch1* mutant showed a delayed induction of the LtD NPQ response, requiring an extra 15 s to exhibit LtD NPQ following the light-to-dark transition (**Fig. 4b**, shifted red arrow).

Next, we tested higher order mutants lacking the minor LHCIIs (*NoM*, Δ*LHCB4-6*). The LtD NPQ phenotype was masked or absent in *NoM* leaves with functional but kinetically distinct NPQ relative to WT (**Fig. 1b**, **Fig. 4c**). However, an LtD NPQ response was still present in higher order *NoM* mutants lacking qE (*NoM npq4*) or qE and qZ (*NoM npq1 lut2*), in which most of the quenching and relaxation capacity *in vivo* is abolished (**Fig. 4c**). Unexpectedly, all *NoM* lines showed a weak NPQ induction phenotype in the dark for the entire duration of both 10 min dark cycles (**Fig. 4c**). Despite this sustained dark-induced NPQ phenotype, *NoM* lines relaxed to significantly longer fluorescence lifetimes and lower residual NPQτ than WT in the presence and absence of qE/qZ (**Fig. 4d**).

### LtD NPQ is affected by photoinhibition

Having shown that LtD NPQ is still present in the absence of qE, qZ, qT, qH, as well as the major and minor LHCII, we next interrogated possible contributions related to photoinhibition, or qI. However, lincomycin, an inhibitor of PSII repair, only had a negligible effect on NPQτ kinetics, increasing the amount of residual NPQ in WT leaves (**Supp.** Fig. 1a) but resulting in no significant differences in *npq4* leaves (**Supp.** Fig 1b) under our short-term actinic light regime. To better resolve the role of qI in LtD NPQ we used a longer-term lincomycin pre-treatment to specifically deplete functional reaction centers (RCs) and remove, rather than increase, contributions of qI, similar to efforts that have been previously described (Belgio et al., 2012; Belgio et al., 2015).

Recognizing the correlation between photoinhibition and light intensity (Tyystjärvi and Aro, 1996), and to best constrain the length of the treatment period, WT and *npq4* leaves in the presence or absence of lincomycin were exposed to high light (400 μmol photons m^−2^ s^−1^) for 16 h and dark-acclimated for 1 h prior to TCSPC measurements. In WT + HL-treated leaves, the ability to sustain steady-state NPQ was significantly reduced after treatment with lincomycin (**Fig. 5a**). Steady-state NPQτ was modestly lower in HL-treated *npq4* leaves compared to *npq4*, with a decline in maximum NPQτ of ∼0.3 (**Fig. 5b**, **Fig. 1b**). Surprisingly, NPQτ was almost entirely abolished in HL- and lincomycin-treated *npq4* leaves (**Fig. 5b**). Closer inspection of the *npq4* + HL + lincomycin data shows that despite the low overall NPQ capacity, there appears to be a loss of detectable LtD NPQ, particularly in the 2^nd^ HL-D cycle, albeit with substantial biological noise at such low NPQτ levels (**Fig. 5c**). Consistent with multi-week lincomycin treatment (Belgio et al., 2012), we observed a specific decline in the RC core protein D1 without substantial differences in other chloroplast proteins such as LHCII and the Rubisco large and small subunits (**Fig. 5d, Supp.** Fig 2a,b). Fluorescence lifetime was significantly reduced after HL + lincomycin treatment, but not in the presence of HL or lincomycin alone across both genotypes (**Fig. 5e**).

**Figure 5.**
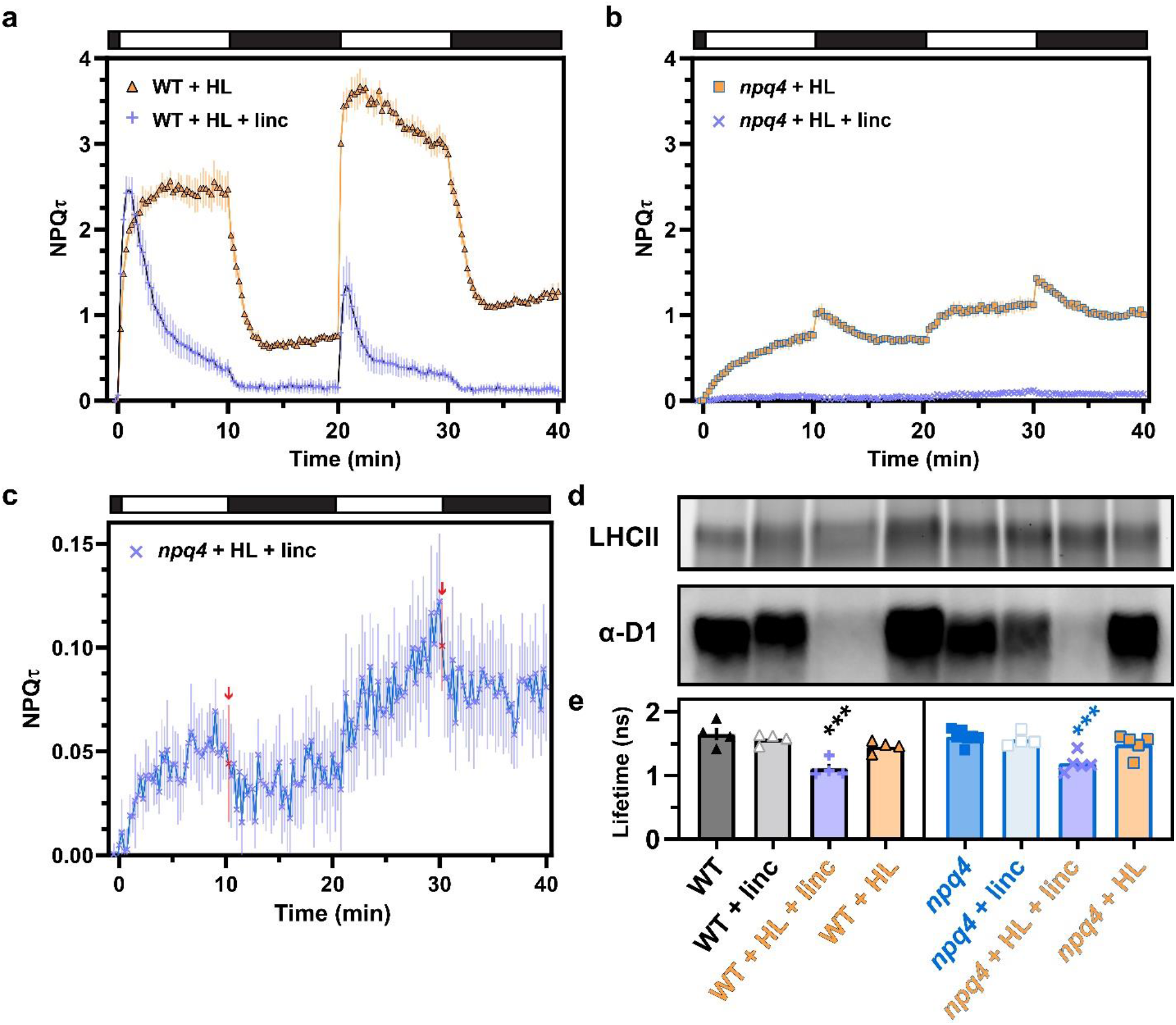
*NPQτ and fluorescence lifetime values of HL-treated WT and npq4 leaves with and without lincomycin*. **(a)** NPQτ of WT pre-treated with HL (black-orange, n=4), and WT pre-treated with HL + 5 mM lincomycin (black- periwinkle, n=4). **(b)** NPQτ of *npq4* pre-treated with HL (blue-orange, n=5), and *npq4* pre-treated with HL + 5 mM lincomycin (blue-periwinkle, n=5). **(c)** Increased NPQτ resolution of *npq4* pre-treated with HL + lincomycin. Red arrows and symbols denote the first time point after the light-dark transition. **(d)** Representative Coomassie stain and immunoblot (1 µg total chlorophyll, 3 biological replicates each) resolving the dominant LHCII band and differences in D1 protein abundance, respectively. Samples are loaded as in (e), with an additional set of replicates shown in *Supplementary* Figure 2. **(e)** Dark-acclimated initial fluorescence lifetime after HL (orange), lincomycin (gray or blue-gray) or HL + lincomycin (periwinkle) treatment for WT (black) and *npq4* (blue) leaves. For all panels error bars show ±1 SEM. Asterisks in (e) indicate significance as determined by ordinary one-way ANOVA using Dunnett’s multiple comparisons test against the untreated genotype (*** p < 0.001).

### Quantification of LtD quenching across treatments and genotypes

The diversity of genotypes and treatments used in this study allowed for quantification and comparison of differences in the magnitude of LtD quenching. To normalize phenotypes across varying fluorescence lifetimes and NPQ capacities, we quantified LtD quenching as the normalized change in fluorescence lifetime (τ) surrounding the light-to-dark transition using the following equation, analogous to the NPQτ calculation:

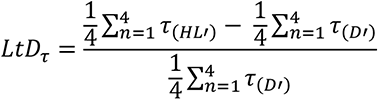

where 𝜏_(𝐻𝐿′)_ and 𝜏_(𝐷′)_ represent the averaged fluorescence lifetimes 1 min before and after the light-to-dark transition, respectively. Thus, the magnitude of LtD_τ_ describes the proportion of LtD NPQ relative to the sum of all NPQ processes, and a higher value equates to a higher proportion of LtD NPQ.

The LtD_τ_ for all higher order *npq4* mutants and treatments are reported in **Fig. 6a**, after correcting for the delayed LtD NPQ phenotype observed in the *soq1 npq4 ch1-4* line (see figure legend). Both HL treatment and loss of chlorophyll *b* resulted in a significantly higher proportion of LtD quenching. In contrast disruptions in ΔΨ via nonactin infiltration, which were roughly comparable in magnitude to the reductions observed by ΔpH disruption by nigericin, significantly reduced the proportion of LtD_τ_.

**Figure 6.**
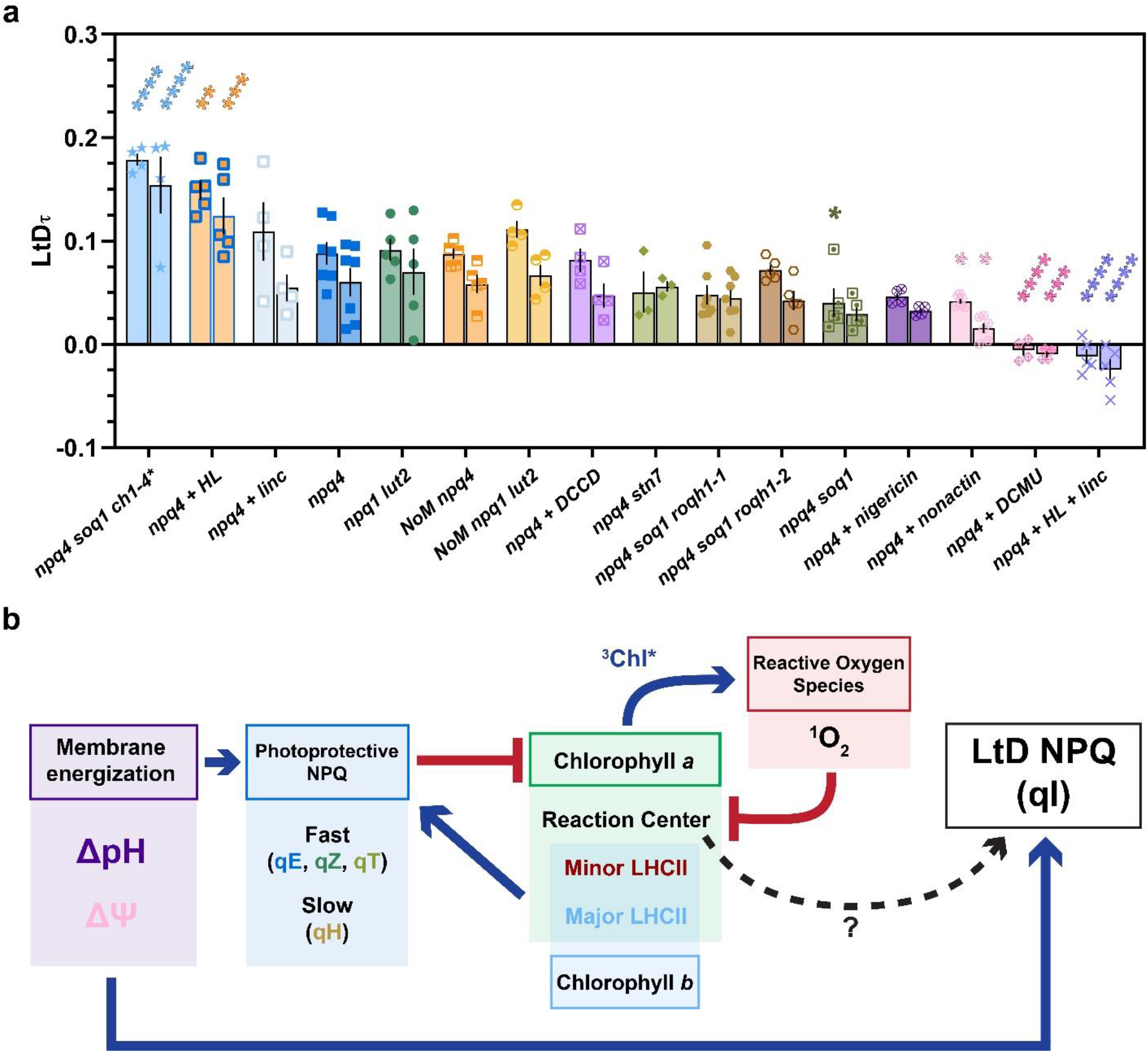
*LtD_τ_ quenching dynamics resolved across all tested npq4 genotypes and treatments*. (**a**) Proportion of fluorescence lifetime quenching attributed to LtD NPQ (LtD_τ_) for the first (t = 10 min, left) and second (t = 30 min, right) light-dark transitions. The τ_(D’)_ used to calculate LtD_τ_ for *npq4 soq1 ch1-4* was adjusted to account for the delay in LtD NPQ induction (t = 15 s to 75 s after the light-dark transition, noted by an asterisk *). Conditions resulting in a statistically significantly different LtD_τ_ from *npq4* were determined by two-way ANOVA using Dunnett’s multiple comparisons test against *npq4* (* p < 0.05, ** p < 0.01, *** p < 0.001, **** p < 0.0001). Error bars show ±1 SEM. (**b**) A flowchart summarizing phenotypes reported in Fig. 1-6. Blue arrows indicate contributing factors whereas red lines indicate inhibiting factors to the subsequent flowchart step. Dashed lines indicate hypothesized relationships determined through this work.

Treatment with DCMU entirely abolished detectable LtD NPQ. The only tested condition where the *npq4* line exhibited detectable NPQ relaxation was after combined photoinhibitory HL and lincomycin treatment (p < 0.0001), particularly in the 2^nd^ HL-D cycle, though this treatment also resulted in the loss of a vast majority of detectable NPQ (**Fig. 5c**). A working model summarizing these phenotypes is shown in **Fig. 6b**, and is discussed in detail below.

## DISCUSSION

NPQ mechanisms are essential but complex modulators of photosynthetic efficiency and plant fitness. A finer understanding of this diverse suite of processes is necessary to unravel still unknown details regarding the key sites and molecular players of photoprotection. One such unknown is the presence of a non-canonical, light-to-dark transition-dependent NPQ signature that has been observed in diverse photosynthetic organisms lacking functional qE. Our study utilized two, sequential HL-D cycles to interrogate the existence and magnitude of this LtD quenching behavior in the model organism *Arabidopsis thaliana*. By using various chemical treatments and higher order *npq4* mutants, we unravel the possible origin of LtD NPQ and discuss its implications in our broader understanding of photoprotection and NPQ relaxation.

First, we used chemical inhibitors to resolve how membrane energetics affects the formation of LtD NPQ. Contrary to our expectations, infiltration of DCCD, which binds protonatable residues of lumen-exposed proteins like PsbS under acidic conditions (Li et al., 2004), did not competitively inhibit PsbS as reported in isolated thylakoids (Ruban et al., 1992). However, DCCD also binds essential protonatable residues of proteins such as ATP synthase (Jahns and Junge, 1989), whose inhibition likely prevents the functional utilization of ᐃpH and resulted in sustained WT qE in the dark (**Fig. 2a**). While DCCD had no significant effect on NPQ in the *npq4* mutant, collapsing the ΔpH via nigericin or ΔΨ via nonactin reduced the apparent formation of LtD NPQ (**Fig. 2d**). Treatment with nigericin resulted in decreases in LtD_τ_ below the threshold of significance (**Fig. 6a**), although this may be due in part to the sustained quenching in the dark observed in *npq4* lines treated with membrane ion-gradient uncouplers (**Fig. 2c**). The contribution of the electrochemical gradient is further supported by complete inhibition of LtD NPQ by DCMU, suggesting an essential role for linear electron transport and/or the Q_B_ site within the PSII RC. Altogether, these data suggest that the lumen acidification, proton motive force, and/or ion gradient across the thylakoid membrane are important contributors to both WT NPQ and the LtD phenotype (**Fig. 2a**, 2b).

The use of various Arabidopsis mutants further expanded our quantitative understanding of LtD NPQ. Higher order *npq4* mutants deficient in qZ, qT, and the carotenoids Zea and Lut still exhibited LtD NPQ (**Fig 3a**). Lines with constitutive qH still had LtD NPQ, even though overall NPQ capacity was attenuated (**Fig. 3b**). The presence of LtD NPQ in these lines aligns with previously observed LtD NPQ phenotypes in *npq4 lcnp* knockout mutants that lack qH altogether (Malnoë et al., 2018). Additionally, loss of the minor LHCIIs (*NoM*) or the loss of chlorophyll *b* (*ch1*) and concomitant loss of LHCII protein stability (Murray and Kohorn, 1991; Espineda et al., 1999) did not abolish the LtD NPQ phenotype (**Fig. 4a**, 4c), despite significantly affecting the overall magnitude and kinetics of NPQ. These results imply the existence of a reaction center-associated NPQ mechanism independent of Zea and Lut that is observable after light-to-dark transitions.

To test this, we used pre-treatments of HL and/or lincomycin, a chemical inhibitor of plastid translation and PSII repair. Interestingly, we observed an increased LtD NPQ response in *npq4* leaves acclimated to HL alone (**Fig. 6a**). We found no significant difference in initial fluorescence lifetime between untreated and HL-treated plants (**Fig. 5e**), suggesting photodamage or constitutive quenching was not responsible for the observed increased LtD_τ_ phenotype. Alternate explanations may include increases in the Chl *a/b* ratio with HL-dependent reductions in LHCII antenna size, which are consistent with the increases in LtD NPQ observed in the *npq4 soq1 ch1* mutant that lacks chlorophyll *b* and most LHCII (**Fig. 6a**). Both conditions would increase the RC:antenna ratio, possibly providing fewer sites for energy trapping and photoprotective NPQ in the PSII supercomplex. In contrast, the inverse of these conditions (a significant decrease in the RC:antenna ratio in HL- and lincomycin-treated *npq4* leaves) abolished almost all NPQ, including the contributions of LtD NPQ (**Fig. 5c**, **Fig. 6a**), emphasizing the importance of PsbS and RCs in both rapid photoprotection and LtD NPQ.

Surprisingly, we also see a loss of the ability to sustain a robust NPQ response in WT leaves depleted of D1 and functional PSII RCs (**Fig. 5a**, HL- and lincomycin-treated WT), which is the opposite of the robust NPQ reported in isolated chloroplasts from plants with varying PsbS protein abundances similarly depleted of their PSII (and PSI) RCs (Ware et al., 2015). Our results, based on fluorescence lifetime snapshots of whole leaves after shorter lincomycin-treatment windows with D1 depletion by HL, suggest that the quenching state produced by PsbS and isolated or aggregated LHCII *in vivo* is transient, and that sustained WT NPQ is dependent on PSII-LHCII supercomplex connectivity.

Returning to LtD NPQ, we propose two hypotheses that explain its existence and relevance to photosynthesis. The first hypothesis suggests that LtD NPQ always occurs upon light-to-dark transition, but our ability to resolve it is masked by the large magnitude of qE relaxation in WT. This is consistent with the reduced magnitude of LtD NPQ when partially masked by *soq1*-dependent NPQ relaxation (*npq4 soq1*, **Fig. 4a,b**). While this evidence is weakly supported by reductions in the proportion of LtD_τ_ in the first light-dark transition for *npq4 soq1* (p < 0.05, **Fig. 6a**), it is also possible that qH is induced and relaxed independently from LtD NPQ. This would also explain why LtD NPQ was still present despite constitutive qH-type quenching of LHCII trimers (Bru et al., 2022) in the *npq4 soq1 roqh1* mutants (**Fig. 3b**, 6a). Furthermore, the absolute change in *npq4* fluorescence lifetime at the first 15 s timepoint after each light-to-dark transition is also large relative to WT fluorescence lifetime recovery (**Fig. 1c**), suggesting that LtD NPQ in WT would at least have to be diminished or suppressed, if present.

Our second hypothesis suggests LtD NPQ is related to a type of photoinhibition mitigated by photoprotective NPQ coordinated by the LHCII antenna. The rapid, transient nature of this phenomenon could be explained by high light-induced changes in membrane energization (i.e. ΔpH, ΔΨ), which may affect (increase) the fluorescence lifetime of the qI state in the *npq4* mutant. Upon light-to-dark transitions, the membrane energization relaxes and its effect on the lifetime of the qI state goes away, resulting in a small, sudden decrease in lifetime (∼0.05 ns, **Fig. 1c**) and corresponding increase in quenching proportional to the amount of qI induced during each light period. This is consistent with increases in LtD_τ_ in the higher order *ch1* mutant, which is also known to have significantly higher production of photoinhibitory singlet oxygen (^1^O_2_) in the light (Dall’Osto et al., 2010), and is further substantiated by the loss of the LtD quenching state after treatment with DCMU, which inhibits both membrane energization and ^1^O_2_ production (Fischer et al., 2006) (**Fig. 2a, 2b**). The reason for the delayed onset of LtD NPQ in the *npq4 soq1 ch1* mutant remains ambiguous, but loss of chlorophyll *b* has been reported to differentially affect lipid and LHCII mobility in the thylakoid membrane (Tyutereva et al., 2017) which may also affect the rate of formation of the qI quenching state.

Our process-of-elimination approach suggests unprotected chlorophyll *a* within the PSII RCs are the likely contributors of LtD NPQ *in vivo* (**Fig. 6b**). The question remains whether this putative Chl-RC quenching is photoinhibitory (qI) or a distinct form of NPQ. There is ample evidence that the RC may serve as a site of NPQ, though at a reduced magnitude relative to quenching facilitated by associated LHCII proteins (Nicol et al., 2019). Our results are also consistent with analysis of hours-timescale qI in the alga *Chlamydomonas reinhardtii*, where a photo-oxidized Chl on the D1 RC protein was proposed as an essential site for qI quenching *in vivo* (Nawrocki et al., 2021). Our HL- and lincomycin-treatments depleted the RC core protein D1 and thus would have functionally removed the detectable contributions of this Chl-D1 site (**Fig. 5d**). This would explain the lack of LtD NPQ and qI in npq4 + HL + lincomycin treated leaves (**Fig. 5b**).

While RC quenching in photo-inactivated PSII cores has historically been attributed to charge recombination between PheoQ_A_^-^ and P680^+^ (Vass et al., 1992), others have proposed the existence of other possible RC quenching thermal traps (Napiwotzki et al., 1997). For example, transient increases in NPQ during transitions from dark to sub-saturating light (Finazzi et al., 2004), and the existence of two distinct pools of photo-inactivated PSII with varying quenched fluorescence lifetimes (Matsubara and Chow, 2004) support a mechanism in which photo-inactivated PSII serve as strong functional quenchers at the RC level that may protect the dwindling pool of functional RCs necessary to sustain ATP synthesis and PSII repair. Such a mechanism may have been an ancestral form of NPQ (Sun et al., 2006; Ivanov et al., 2008), or an artifact from cyanobacteria that employ mechanisms such as LHC-like high-light inducible proteins (HLIPs) to quench PSII intermediates and suppress photo-oxidative damage (Staleva et al., 2015).

Based on these cumulative findings, we find strong evidence that the evolutionarily conserved, non-canonical LtD NPQ signature is related to qI quenching that can be observed in the absence of qE and is exacerbated in RCs that lack photoprotective NPQ. We hypothesize that this form of quickly induced, but slowly relaxing photoinhibition shares the same fundamental basis as the hours-timescale qI-type photoinhibition found in WT contexts and reveals an effect of membrane energization on the qI state.

This shared mechanism may also explain the greater amounts of residual NPQ that have been previously and consistently observed in lines lacking qE following HL exposure (**Fig. 1b**), indicative of photoinhibition. Importantly, qI is predicted to have significant, competitive costs to photosynthetic efficiency and biomass accumulation (Long et al., 1994; Murchie and Niyogi, 2011). Additional biochemical, spectroscopic, and mechanistic studies may support efforts to investigate and engineer photoprotective mechanisms that suppress qI without inhibiting overall productivity. Future work assessing the membrane-level dynamics of this process will aid in our understanding of NPQ under fluctuating and photoinhibitory light conditions.

## Acknowledgments

We thank Alizée Malnoë for kindly sharing *npq4 soq1* seeds, Raquel Ponce for her assistance in genotyping the *npq4 soq1 ch1-4* line, and Masakazu Iwai for his insights. This work was supported by the U.S. Department of Energy, Office of Science, Basic Energy Sciences, Chemical Sciences, Geosciences and Biosciences Department. D.P.-T. was supported by the Berkeley Fellowship and the NSF Graduate Research Fellowship Program (Grant DGE 1752814). K.K.N. is an investigator of the Howard Hughes Medical Institute. This article is subject to HHMI’s Open Access to Publications policy. HHMI lab heads have previously granted a nonexclusive CC BY 4.0 license to the public and a sublicensable license to HHMI in their research articles. Pursuant to those licenses, the author-accepted manuscript of this article can be made freely available under a CC BY 4.0 license immediately upon publication.

## Author Contribution

L.L. and D.P.-T. contributed equally. L.L., D.P.-T., G.R.F., and K.K.N. designed the experiment. D.P.-T. selected and screened genotypes and treatments for analysis, with support from A.L. and S.A.M. L.L. performed infiltrations and the TCSPC measurements. D.P.-T. and L.L. analyzed the data with input from H.E.L., C.J.S., A.M., and T.-Y.L. D.P.-T. and L.L. drafted the manuscript with input from G.R.F., K.K.N., and C.J.S.

---

**Supplementary Figure 1.**
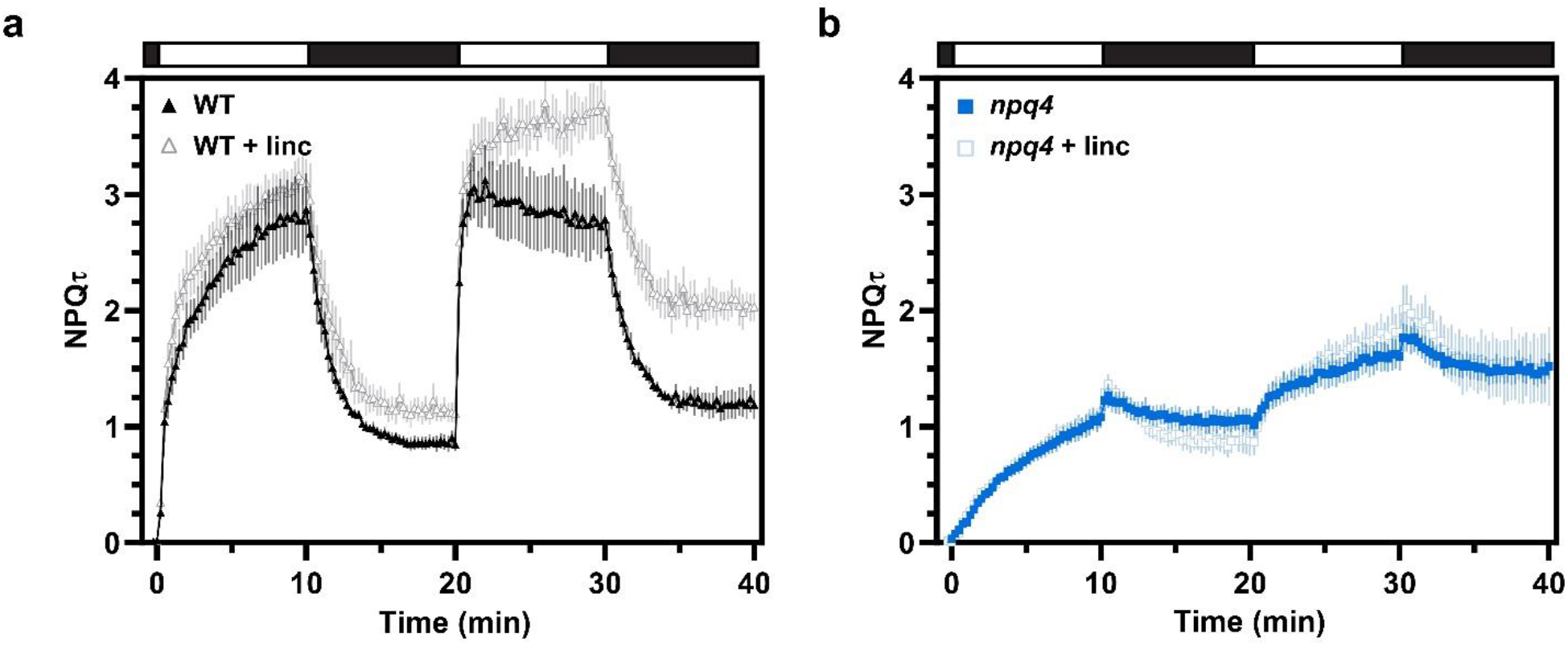
NPQτ of WT and npq4 leaves with and without lincomycin. (a) NPQτ of WT (black, n=4) and WT leaves infiltrated with 5 mM lincomycin (gray, n=4). (b) NPQτ of *npq4* (blue, n=7) and *npq4* leaves infiltrated with 5 mM lincomycin (blue-gray, n=4). Error bars show ±1 SEM.

**Supplementary Figure 2.**
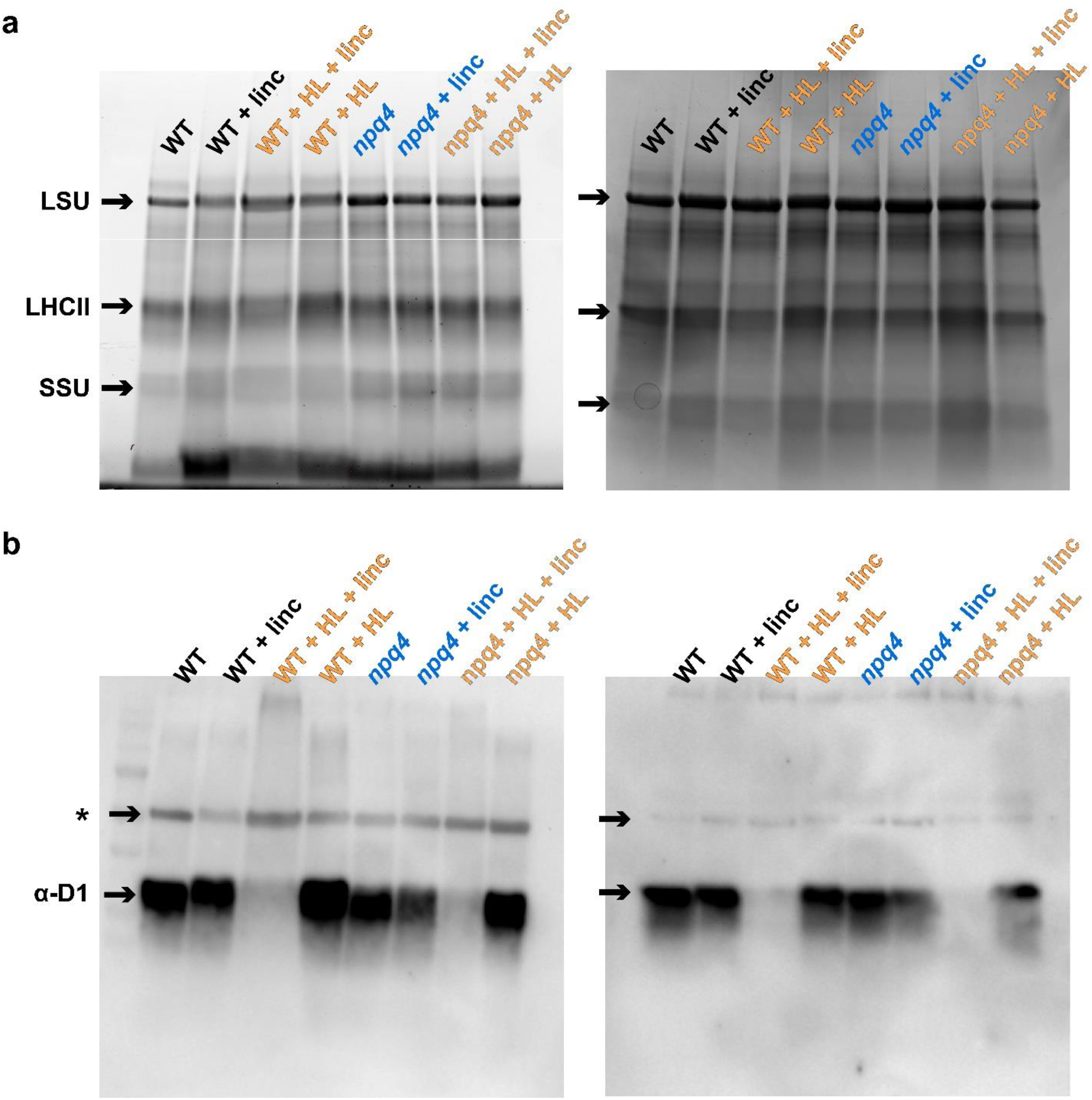
Protein abundances for WT and npq4 leaves treated with HL and/or lincomycin. (a) Coomassie stains showing relative total protein profiles with dominant bands for the rubisco large subunit (LSU), monomeric LHCII, and rubisco small subunit (SSU) denoted by arrows. (b) Immunoblot detection for differences in D1 (PsbA) protein abundance across genotypes and treatments. A non-specific background band is denoted by an asterisk. For all blots, 1 ug total chlorophyll was loaded in each well, representative of 3 biological replicates.

